# An Alternative Approach to Compute the Likelihood for the DAISIE (Dynamic Assembly of Island biota through Speciation, Immigration, and Extinction) Model

**DOI:** 10.1101/2025.09.03.673763

**Authors:** Ornela N. Dehayem, Bart Haegeman, Rampal S. Etienne

## Abstract

Understanding island biodiversity requires studying the processes driving its assembly and examining how these processes vary over time, across lineages, and across different islands. The DAISIE (Dynamic Assembly of Island biota through Speciation, Immigration, and Extinction) framework has been developed for this purpose, and has advanced our understanding of island community assembly by estimating, from phylogenetic data, the contribution of the processes of colonization, speciation and extinction to island community assembly. However, the model assumes uniform colonization and diversification rates across lineages, ignoring potential variation in these rates caused by, for example, lineage-specific traits. This assumption thus restricts the model’s capacity to capture complex colonization and diversification dynamics, and may consequently bias inference of colonization and diversification dynamics on islands. An extension of the framework behind these more complex dynamics is therefore desired. However, for this state-dependent colonization and diversification extension, the current computation of the likelihood of the model given phylogenetic data is computationally prohibitive. In this study, we therefore propose an alternative approach to computing the DAISIE likelihood, under the assumption of diversity-independent colonization and diversification. Our novel approach is based on the pruning algorithm (which calculates the likelihoods of comparative data given a tree and a model) that has been used for computing the likelihood of SSE (State-dependent Speciation and Extinction) models, which traces lineage history backward in time from the present day to the root of the tree. We demonstrate that our alternative approach reproduces DAISIE’s predictions under diversity-independent colonization and diversification. This is important because no independent evaluation of the likelihood computation in DAISIE has been available. Our alternative approach not only confirms DAISIE results for diversity-independent colonization and diversification, it also gives more confidence in the results under diversity-dependence. Furthermore, our alternative approach offers greater computational efficiency. Finally, it provides a flexible framework for incorporating state-dependent dynamics in colonization, speciation, and extinction processes.

## 1 Introduction

Biodiversity, the variety of life forms on Earth, is a fundamental aspect of the natural world that underpins ecosystem function and resilience (O’Connor et al., 2021; Yadav et al., 2023). Understanding the processes that create and sustain biodiversity is a central goal in ecology and evolution. One of the primary approaches to studying the accumulation of biodiversity over evolutionary time scales is to reconstruct the past dynamics of the two evolutionary processes, speciation and extinction, that are thought to shape species richness over these time scales (Morlon, 2014; Morlon et al., 2022). Diversification models fitted to time-calibrated phylogenies of extant species have been widely used to test evolutionary scenarios that may have shaped patterns of diversification through time (Condamine et al., 2013; Etienne et al., 2012; Etienne and Rosindell, 2012; Maddison et al., 2007; Morlon, 2014; Nee et al., 1994). Each model encodes a distinct scenario of speciation and extinction dynamics − either constant-rate, diversity-dependent, time-dependent, trait-dependent, or assuming protracted speciation. These hypotheses are evaluated by fitting a range of candidate models to identify the scenario that best approximates diversification patterns.

Marine islands present ideal systems to unravel evolutionary processes that have shaped present-day life due to their isolation, well-defined boundaries, and often exceptionally high biodiversity (Gillespie and Roderick, 2014; Helmus et al., 2014; Losos and Ricklefs, 2009; Patino et al., 2017; Warren et al., 2015). Their relatively recent geological origins and limited species richness compared to the much larger continental regions make studying processes shaping biodiversity more tractable. Moreover, islands (in an archipelago or across the globe) offer repeated natural experiments, providing multiple case studies for testing and contrasting evolutionary patterns across diverse environments, time and lineages (Losos and Ricklefs, 2009). Island communities assemble through in situ speciation and extinction as well as via colonization (MacArthur and Wilson, 2001). The DAISIE model (Dynamic Assembly of Island biota through Speciation, Immigration, and Extinction) has been developed to incorporate these three fundamental processes driving island diversity (Etienne et al., 2023; Valente et al., 2015). DAISIE makes use of information contained in phylogenies to assess the role of the processes of colonization (arrival and establishment of new island species from a nearby continent), speciation via anagenesis (evolution to an island species distinct from the mainland ancestor due to reduced gene flow between island and mainland populations) and cladogenesis (the process of an ancestral species giving rise to two different species on the island or archipelago), and extinction (loss of species from the island) in shaping island communities. It also assesses whether the rates of colonization and cladogenesis depend negatively on island diversity, and hence limit diversity.

The DAISIE framework has been successfully applied to 1) quantify how colonization, speciation, and extinction rates vary with island age, area, and isolation (Valente et al., 2020); 2) assess the role of diversity limits and equilibrium processes in shaping patterns of species richness of birds in the Galápagos and Macaronesian islands (Valente et al., 2017, 2015); 3) estimate the amount of evolutionary time lost or threatened due to anthropogenic extinctions (Valente et al., 2017, 2019); 4) demonstrate how environmental changes, such as lake expansion, can elevate equilibrium diversity by enhancing colonization rates (Hauffe et al., 2020), and 5) study the phylogenetic limits to diversity-dependence (Etienne et al., 2023).

While the DAISIE framework has been instrumental in advancing our understanding of community assembly on islands, the current DAISIE model for which parameter estimation is possible through maximum likelihood has certain limitations. The likelihood computation currently requires uniform diversification and extinction parameters across lineages, thus ignoring variability associated with species-specific traits. However, species differ in dispersal abilities, physiological and morphological traits, habitat preferences and ecological interactions, all of which can modulate colonization frequency, speciation speed, and extinction risk. Ignoring such trait dependence can bias model inferences, especially if species’ traits shape the structure of phylogenetic trees on islands (Maddison, 2006; Maddison et al., 2007; Xie et al., 2023). Although DAISIE has been shown to effectively reconstruct biodiversity dynamics in many trait-dependent scenarios, it fails in some scenarios, notably when applied to phylogenetic data of community with large clade size variation (Xie et al., 2023). This limitation highlights the need for a trait-explicit modeling framework. However, extending the DAISIE framework to account for trait dependence is not straightforward, as the underlying mathematical formulation is not readily designed to incorporate trait-specific diversification and colonization dynamics. In this study, we introduce an alternative methodology for modeling community assembly in isolated ecosystems under the assumption of diversity-independent diversification. Our approach follows the SSE (state-dependent speciation and extinction) framework introduced by Maddison et al. (2007), which operates backward in time, tracing lineage histories from the present day back to the island’s origin. We demonstrate that our method aligns with DAISIE’s predictions, while improving computational efficiency by making optimal use of the assumption on diversity-independence. Moreover, it provides a flexible framework that allows the incorporation of state-dependent colonization, speciation, and extinction processes.

## 2 Model and likelihood computation

### 2.1 Data description and assumptions

Our new methodology consists of an alternative approach to compute the likelihood of the DAISIE model as laid out in Valente et al. (2015). The DAISIE model assumes that after the emergence of an island, species can immigrate to the empty island from a mainland with a fixed species richness. Each colonization event initiates an independent lineage that may lead to 1) a single nonendemic species (same as the mainland ancestor), 2) a singleton endemic species (via anagenesis or cladogenesis followed by extinction), 3) a radiation of endemic species (via cladogenesis), or 4) extinction without leaving any extant descendants. The island community is thus represented by one or more such lineages, each corresponding to a separate colonization event and represented by its own phylogenetic tree. Fig. 1 provides an example of a dataset for an island community. Each clade represents a separate colonization event from the mainland that has led to one or more species on the island. The dots represent the estimates of the colonization times, which are inferred by measuring the genetic divergence between island species and their mainland sister species (or mainland populations of the same species in the case of a non-endemic species). For the cases where these colonization times are unknown, the model integrates over all possible colonization times, that is, between the island-archipelago age and the present for singleton lineages (e.g., C and F), and between the island-archipelago age and the crown age for an endemic clade (e.g., A, B, D, E).

**Figure 1.**
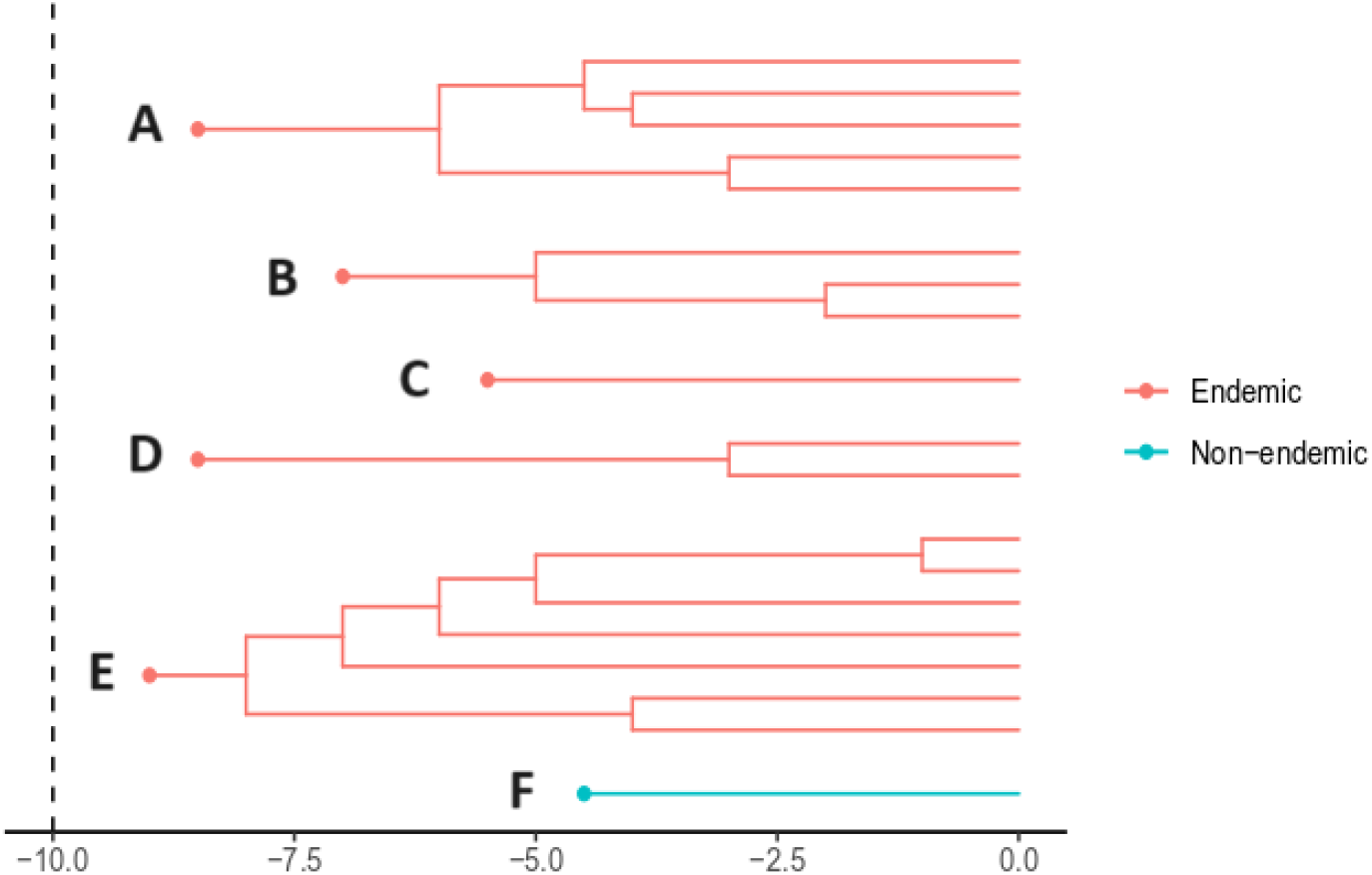
Example of a dataset in the DAISIE framework. Each tree represents a colonization event with one or more descendants on the island. Circles indicate the time of colonization of each lineage; red - endemic lineages; blue - non-endemic lineages. The clades *A, B, D*, and *E* illustrate cases where a mainland species colonizes the island and subsequently gives rise to multiple descendant species. *C* corresponds to a scenario where a mainland species colonizes the island and speciates via anagenesis or cladogenesis, but only a single species survives until the present. *F* represents a scenario where a mainland species colonizes the island and does not undergo speciation.

It is possible that before an observed colonization event, the same mainland species already colonized the island (Caujapé-Castells et al., 2017) (and persisted until the observed colonization event), but the model assumes that the observed lineage descends from the latest colonization event. This means that at the observed colonization event, any trace of these previous colonizations is erased (Valente et al., 2015). Recolonization of the same mainland ancestors can occur and still remain on the island, but only after the previous colonization has been followed by speciation. For this reason, the likelihood calculation assumes recolonization by the same mainland species is only possible after the island species undergoes speciation, because if this happened before speciation, the recolonizing population would have replaced the entire island population, and the most recent colonization event would be considered the new divergence time from the mainland which would be in disagreement with the data. Here we follow the same assumption; we do not question its validity in the current paper but will leave this for a future exploration.

DAISIE can accurately estimate colonization and diversification rates within island communities in most cases from colonization times of independent lineages, their branching times, endemicity status (derived from phylogenetic data), and the geological age of the island (Lambert et al., 2022; Neves et al., 2021; Xie et al., 2023). The model also allows for diversity-dependence in colonization and cladogenetic speciation to act either within clades (clade-specific, CS model) (Valente et al., 2015) or across the entire island community (island-wide, IW model) (Etienne et al., 2023). We note that for our methodology to work we assume that, in contrast to DAISIE, there is no diversity-dependent colonization or cladogenesis. Another key difference lies in how the two models handle incomplete data sampling. DAISIE uses *n*-sampling (Etienne et al., 2012), which assumes that, given a phylogenetic (or reconstructed) tree with a known number of extant species *S*, exactly *n* species are not sampled. In contrast, our approach uses the ρ-sampling scheme (FitzJohn et al., 2009), where the phylogeny is assumed to be a random sample of all extant species, where each species is included in the phylogeny with probability ρ, and with probability 1 − ρ to be unsampled. The two types of sampling are related by

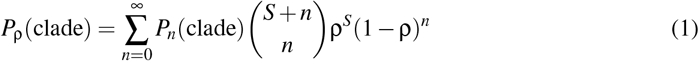

where *S* is the number of species in the clade. In practice, the following procedure would be fol-lowed. Suppose we have a phylogeny with *S* tips and we know that *m* species are not represented in the phylogeny. With *n* sampling we would set *n* equal to *m*, whereas with ρ-sampling ρ is set equal to 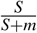.

### 2.2 Likelihood computation

Mainland species are considered independent of one another, allowing the dynamics of each colonizing species to be modeled separately (CS model, see above). The likelihood of observing the entire community is then calculated as the product of the probabilities of each independent colonization plus its potential descendants (which we will refer to as “lineage” hereafter even if it is a non-endemic singleton) present on the island, and accounting for potential colonization events that occurred before and after the observed colonization, and subsequently went extinct. The model distinguishes three types of lineage:

- No surviving lineage: a colonization event that resulted in a lineage that has gone completely extinct and is not represented among the present-day species on the island.
- Non-endemic singleton: a species present on the island that originated from a colonization event and has not undergone speciation since arrival, and is also found on the mainland.
- Endemic singleton: a species present on the island that originated from a colonization event and has undergone anagenetic or cladogenetic speciation, but only one species survives until the present on the island.
- Endemic radiation: a clade of two or more species that all descended from a single colonization event and underwent cladogenetic speciation on the island, resulting in a group of endemic species.

To compute the probabilities of observing a clade, DAISIE uses a Hidden Markov model, based on the variable 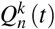 which describes the probabilities that the diversification process is consistent with the observed phylogeny from a starting time to a time *t*, with *k* extant lineages and *n* species present in the phylogeny at time *t*. The probabilities 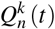 are computed forward in time, from the initial time *t*_0_ at the island origin to the present time *t*_*p*_. In contrast to the original DAISIE model, which traces lineage history forward in time from the island’s origin to the present, our approach traces it backward, from the present day to the island’s origin. We define the backward time τ, measured from the present to the past. It is related to the usual forward time *t* by the equation τ = τ_*p*_ − *t* + *t*_*p*_, where τ_*p*_ is the backward time at the present. We follow the differential equation-based approach introduced by Maddison et al. (2007), where probabilities are assigned at the tips of the lineage, and a set of rules is used to iteratively update these values as we trace the lineage backward in time, from the tips to the colonization time. At the colonization time of a lineage, we incorporate the probability of colonization, before continuing the computation further back in time to the island’s origin.

We define two central probabilities used in the model:

- *E*(τ), the probability that a lineage starting at time τ as an endemic species leaves no descendants at the present;
- *D*(τ), the probability that a lineage alive at time τ leaves a clade that is consistent with the observed data. In order to capture the state of the system at specific points in time, we introduce several state-dependent variants of the probability *D*(τ). These states represent the probabilities of the dynamics being consistent with the observed data. We define these states as follows:
  - *D*_*E*_ (τ) is the probability that a lineage that started on the island at time τ as an endemic lineage evolves into the observed clade;
  - *D*_*M*_(τ) is the probability that all lineages descending from the mainland species present at time τ, as well as those originating from subsequent colonizations, evolve in a manner consistent with the observed data;
  - *D*_*A*_(τ) is the probability that the mainland species is absent from the island at time τ, and that all lineages resulting from subsequent colonizations evolve in a manner consistent with the observed data;

Fig. 2 provides a schematic representation of the state probabilities *D*_*M*_(τ) and *D*_*A*_(τ) defined in the model.

**Figure 2.**
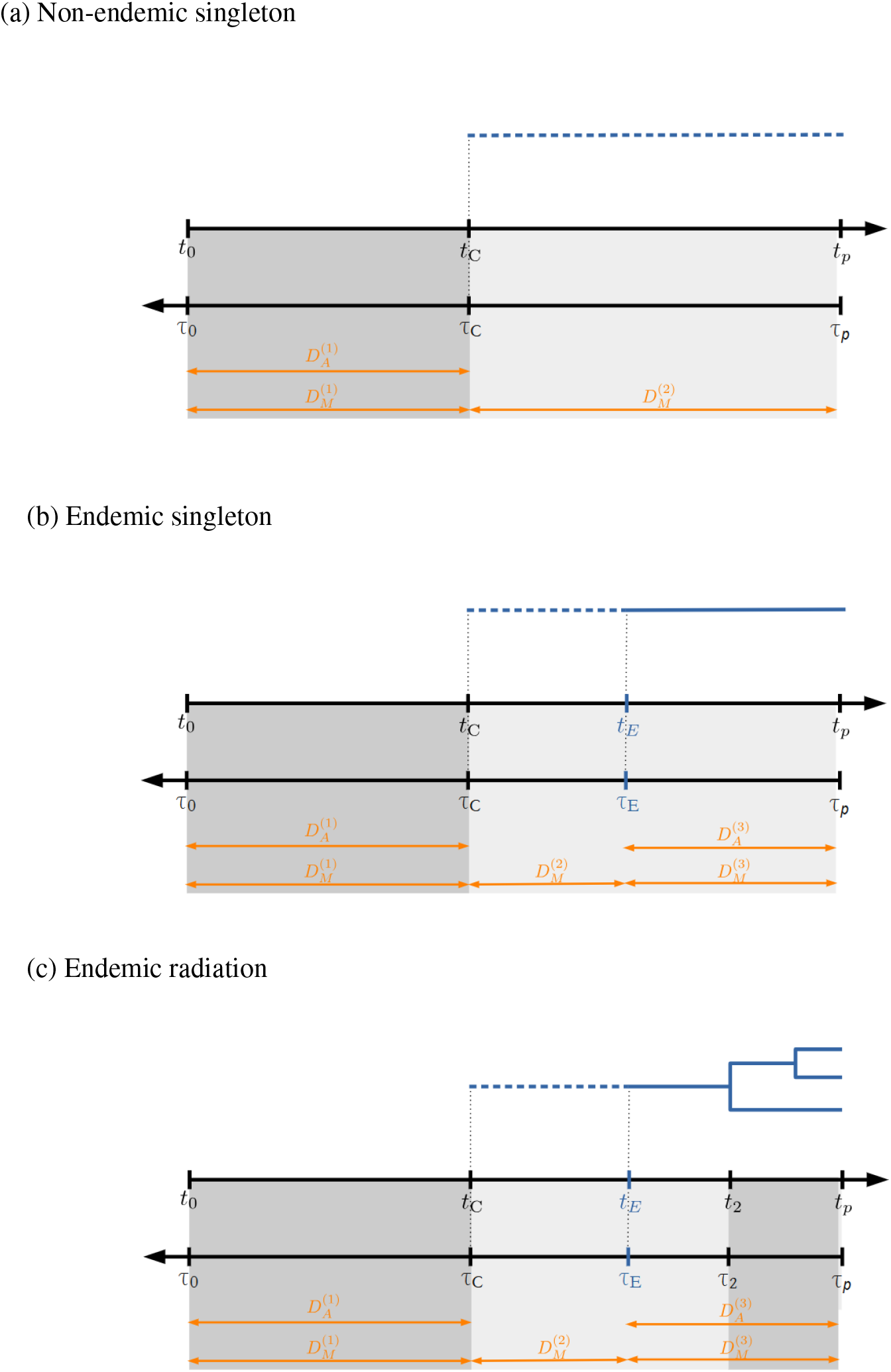
Schematic representations of the temporal structure and definition of the state probabilities *D*_*M*_(τ) and *D*_*A*_(τ) in the backward pruning algorithm for different cases. The dashed blue line represents the time interval during which the species is non-endemic. The blue solid line represents the time interval during which the species is endemic. The orange arrows show the time intervals over which each probability is computed, going backward in time. The grey-shaded regions represent different time intervals, each governed by a distinct set of differential equations. These intervals correspond to different phases in the lineage history, reflecting changes in model dynamics across time.

The times shown in black (τ_0_, τ_*C*_, τ_2_, and τ_*p*_) are extracted from the data. In contrast, the time points shown in blue (τ_*E*_) are realization-dependent. These times are described as follows:

- *t*_0_, τ_0_: time of the island’s origin, i.e. the island’s age
- *t*_*C*_, τ_*C*_: time of colonization event that gives rise to lineage observed at the present time
- *t*_*E*_, τ_*E*_ : time at which the endemic lineage at the present time becomes endemic
- *t*_*p*_, τ_*p*_: present day time

We construct the likelihoods by integrating a series of differential equations. To formulate these equations, it is convenient to introduce different instances of the variables *D*_*A*_ and *D*_*M*_:

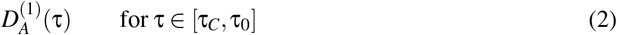

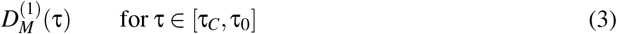

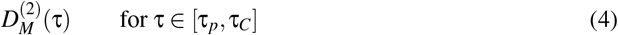

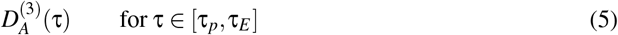

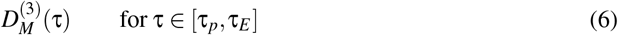

We use the superscripts to distinguish the same variable occurring in different systems of equations.

- 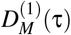, the probability that the dynamics evolves in a clade consistent with the data, starting at time τ from an island with the non-endemic species which is not represented in the clade. This mainland species originates from a colonization event between *t*_0_ and *t*_*C*_. In this state the mainland species goes extinct before the present, or is taken over by the observed mainland species;
- 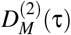, the probability that the dynamics results in a clade consistent with the data, starting at time τ from an island with the non-endemic species which is represented in the clade. This mainland species originates from a colonization at τ_*C*_. In this state the mainland species can speciate to initiate the observed lineage;
- 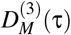, the probability that the dynamics leads to a clade consistent with the data, starting at time τ from an island with the non-endemic species which is not represented in the clade. This mainland species originates from a colonization after the speciation event on the observed lineage at τ_*E*_ . In this state the mainland species goes extinct before the present.
- 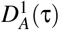 is the probability that the mainland species 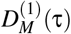 is absent from the island at time τ
- 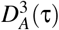 is the probability that the mainland species 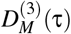 is absent from the island at time τ

Let us consider a lineage present on an island, with estimated colonization time τ_*C*_, which may or may not have diversified into extant species. At the time τ_*C*_ the mainland ancestor colonized the island at a colonization rate γ. Following colonization, the lineage can diversify via cladogenesis at a rate λ^*c*^, or anagenesis at a rate λ^*a*^, or may go extinct at a rate *µ*. Below we describe how to compute the likelihood for different types of lineages with given colonization time τ_*C*_. The equations can be readily extended to allow for a colonization time that is not precisely known but lies between τ_*max*_ and τ_*min*_. These equations are given in the appendix.

#### 2.2.1 No surviving descendants

In this scenario, mainland species may have colonized, but none of the colonists survive to the present τ_*p*_. The dynamics of the mainland species on the island are captured by the pair of vari-ables 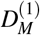 and 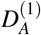. The dynamical equations describing the change in the probabilities from the present time τ_*p*_ to time of the island’s origin τ_0_ are given by:

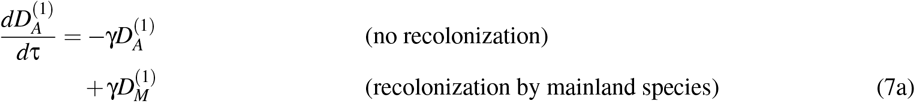

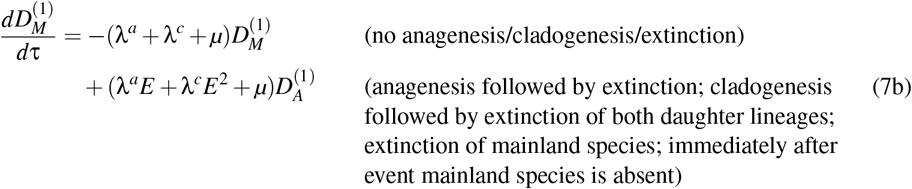

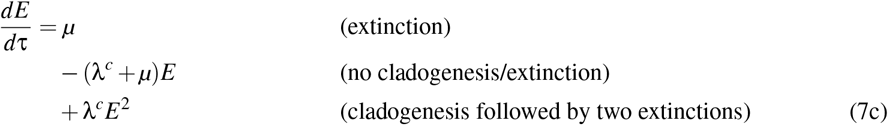

These equations describe the scenario in which lineages colonize the island at some point between τ_*p*_ and τ_0_, may or may not undergo speciation through anagenesis or cladogenesis, but all resulting lineages eventually go extinct before the present.

At the present time τ_*p*_, the initial conditions are:

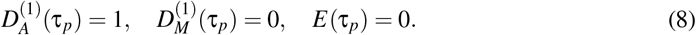

This initialization reflects the fact that the mainland species is not observed on the island at the present time. Integrating these quantities backward to τ_0_ gives the likelihood

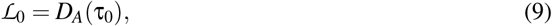

equal to the probability that none of the species colonizing the island between *t*_0_ and the present time *t*_*p*_ leaves any descendants.

#### 2.2.2 Non-endemic singleton lineage, with colonization time at τ_*C*_

In this scenario, the mainland species colonizes the island at time τ_*C*_ and remains non-endemic (no speciation occurs) until the present. In the interval [τ_*p*_, τ_*C*_] the mainland species is present on the island and represented in the clade (which consists of a single branch), so we use the variable 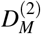. The dynamical equations describing the change in the probabilities between τ_*p*_ and τ_*C*_ are given by:

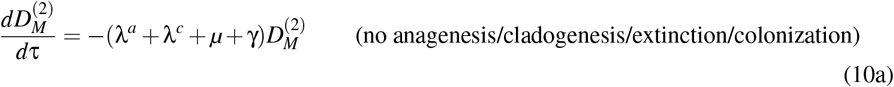

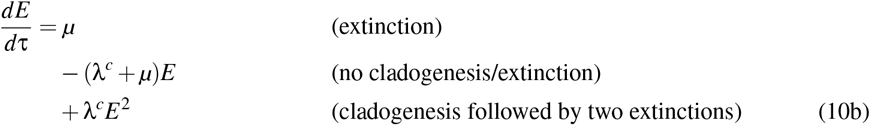

with initial conditions at τ_*p*_ given by:

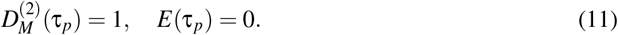

In the interval [τ_*C*_, τ_0_] there is no clade, and so the mainland species is not represented. Still it could have colonized the island, a possibility we track with the variables 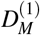 and 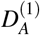. The dynamical equations describing the change in the probabilities between τ_*C*_ and τ_0_ are given by Eqs. (7a−7c), with initial conditions at τ_*C*_ given by:

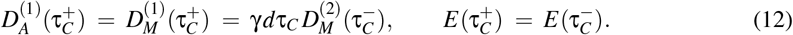

where 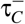 represent the time immediately before τ_*C*_ (“before” in backward time), and 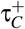 the time immediately after τ_*C*_.

At the island age *t*_0_, the likelihood is

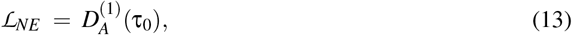

equal to the probability of observing a non-endemic lineage that colonized the island at time *t*_*C*_.

#### 2.2.3 Endemic singleton lineage, with colonization time at τ_*C*_

In this scenario, a mainland species colonizes the island at time τ_*C*_, and speciates at an unknown speciation τ_*E*_ time between τ_*C*_ and τ_*p*_. Thus, between τ_*C*_ and τ_*E*_, the species is present on the island and is non-endemic; its dynamics are captured by the variable 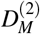. After speciation and before τ_*p*_, the mainland ancestor may recolonize the island and go extinct again; this is captured by the variables 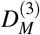 and 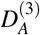. The dynamical equations describing the change in the probabilities between τ_*p*_ and τ_*C*_ are given by:

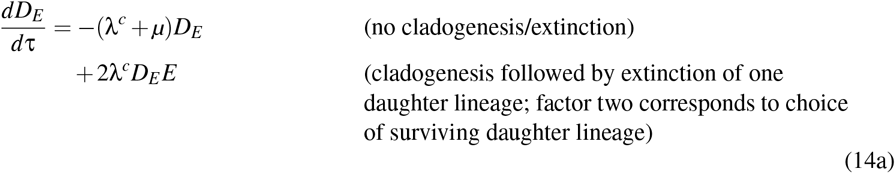

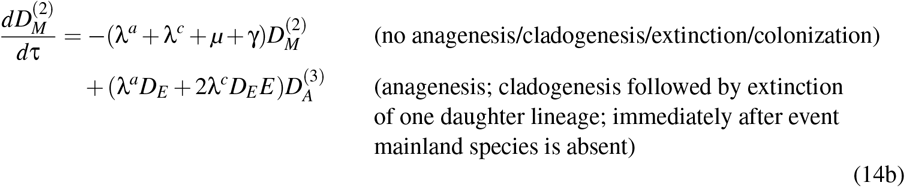

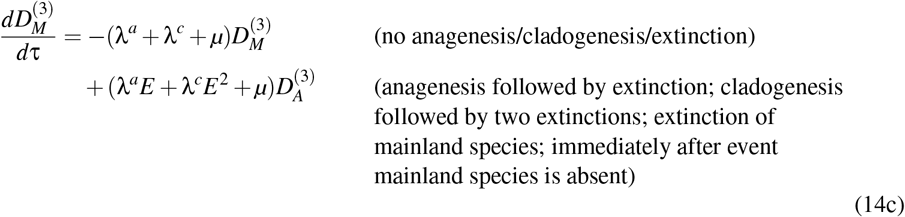

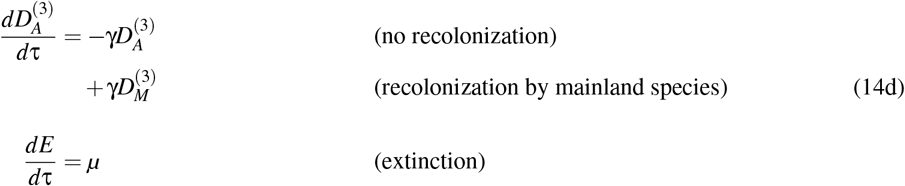

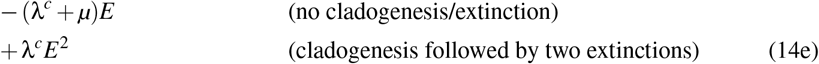

with initial conditions at τ_*p*_ given by:

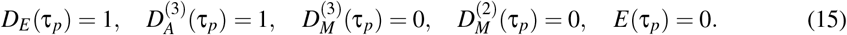

In the interval [τ_*C*_, τ_0_] there is no clade, and so the mainland species is not represented. Still it could have colonized the island, a possibility we describe with the variable 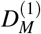. As in the previous case, the dynamical equations describing the change in the probabilities between τ_*C*_ and τ_0_ are given by Eqs. (7a − 7c), with the initial conditions Eq.(12) at τ_*C*_. At the island age τ_0_, the likelihood is

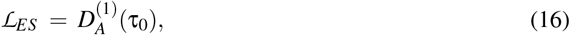

equal to the probability of observing an endemic lineage that colonized the island at time *t*_*C*_.

#### 2.2.4 Endemic radiation, with colonization time at τ_*C*_

In this scenario, the mainland species colonizes the island at time τ_*C*_ and diversifies into several extant descendant species at the present time τ_*p*_. After the first speciation of the lineage, the mainland ancestor may recolonize the island and go extinct again; this is captured by the variables 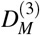 and 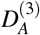. Between the crown age τ_2_ and the present τ_*p*_, the lineage is present, and has already undergone cladogenesis speciation. The dynamical equations describing the change in the probabilities between τ_*p*_ and τ_2_ are given by:

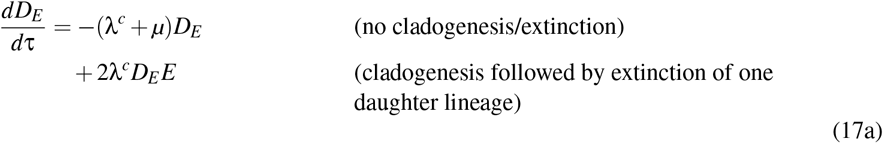

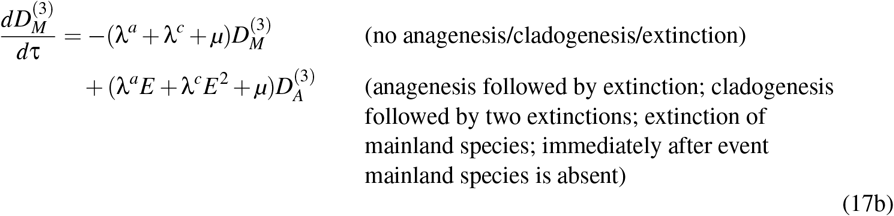

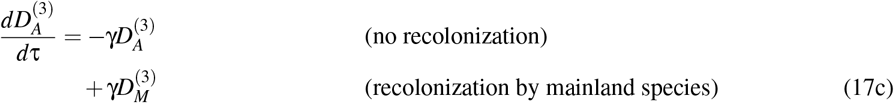

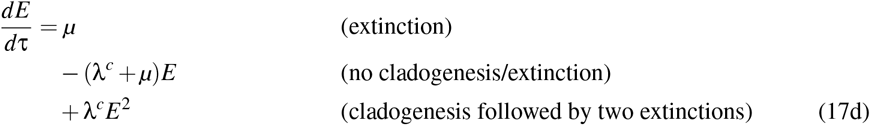

The initial conditions at τ_*p*_ are given by:

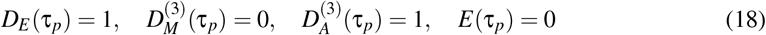

If species in the lineages are sampled at random (all species in the clade have the same probability of being sampled), incomplete sampling can be accounted for by modifying the initial conditions for each tip in the tree to reflect ρ, the probability of sampling a species, following FitzJohn et al. (2009). The initial conditions at τ_*p*_ are modified to:

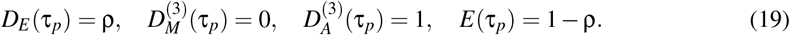

where ρ is the sampling fraction of the lineage.

The set of equations Eqs. (17a − 17d) are integrated along the edges of the trees starting from the tips. At each branching time, the probabilities of the two daughter branches are combined by multiplying both probabilities with one another and with λ^*c*^ dτ which represents the probability of cladogenesis speciation over the small time interval *d*τ. The resulting product serves as the initial condition for the ancestral (subtending) branch as we continue tracing the lineage backward toward its colonization time.

In the time interval [τ_2_, τ_*C*_], the species has already colonized and may or may not have speciated. We have to add the dynamical equation for 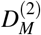, because we know that on at least a part of the branch the species is non-endemic. The dynamical equations describing the change in the probabilities between τ_2_ and τ_*C*_ are given by Eqs. (14a−14e). As initial conditions for 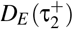 at τ_2_, we take λ^*c*^*d*τ times the product of the *D*_*E*_ functions of the merging subclades. For 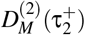 we use the same quantity multiplied by the probability 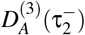 that the mainland species is not present on the island. This covers both cases in which the speciation event that transforms the mainland species into an endemic one occurs either between τ_*C*_ and τ_2_ or precisely at time τ_2_. The initial conditions at τ_2_ are thus:

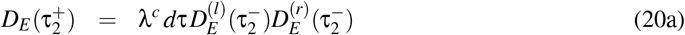

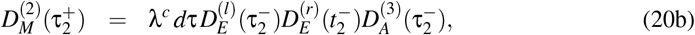

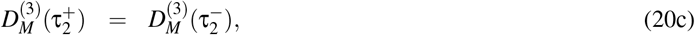

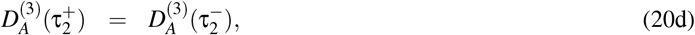

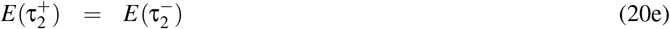

Here 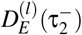 and 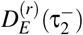 are the probabilities that the two descendant lineages (left and right lineage in the phylogeny) are endemic just before τ_2_.

In the interval [τ_*C*_, τ_0_], there is no clade. The dynamics are the same as in the previous cases. At the island age *t*_0_, the likelihood is

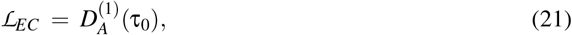

equal to the probability of observing a non-endemic lineage that colonized the island at time *t*_*C*_.

We implemented these likelihood computations, including those for unknown colonization time in the R package DAISIE available on CRAN at https://rdrr.io/cran/DAISIE/, where we refer to them as DAISIE-DE, after the main equations for *D* and *E*.

## 3 Parameter estimates by DAISIE vs DAISIE-DE

To evaluate the performance of our new approach DAISIE-DE, we compared its parameter estimates with those obtained from the DAISIE model across four empirical island biota datasets (Table 1). For each dataset, both models were fitted to estimate the rates of colonization, cladogenesis, anagenesis, and extinction. The carrying capacity was set to *K* = ∞ in DAISIE to eliminate the influence of diversity-dependent dynamics, allowing a comparison of the diversity-independent DAISIE and DAISIE-DE. It is important to note that all datasets used in this comparison were complete, as DAISIE and DAISIE-DE differ in how they account for missing species. Table 1 summarizes the resulting rate estimates. The comparison between the DAISIE and DAISIE-DE results shows that, for each dataset, the maximum likelihood and corresponding parameter estimates of the observed phylogenetic data are practically identical. The minute differences are due to numerical precision limits in integration and optimization.

**Table 1:** Comparison of rates estimated by DAISIE and DAISIE-DE across different empirical datasets.

| Empirical dataset | Method | $\lambda^c$ | $\mu$ | $\gamma$ | $\lambda^a$ | logLik |
| --- | --- | --- | --- | --- | --- | --- |
| Galápagos birds<br>(Valente et al., 2015) | DAISIE | 2.550446 | 2.683555 | 0.009343 | 1.007324 | -75.99241 |
|  | DAISIE-DE | 2.550682 | 2.683818 | 0.009344 | 1.007281 | -75.99241 |
| Hispaniola frogs<br>(Etienne et al., 2023) | DAISIE | 0.178709 | 0.024700 | 0.000626 | 0.000000 | -209.8507 |
|  | DAISIE-DE | 0.178702 | 0.024685 | 0.000626 | 0.000000 | -209.8507 |
| Cape Verde birds<br>(Valente et al., 2019) | DAISIE | 0.000000 | 0.7678714 | 0.026222 | 0.462818 | -62.63102 |
|  | DAISIE-DE | 0.000000 | 0.767607 | 0.026216 | 0.462811 | -62.63102 |
| Biwa fish<br>(Hauffe et al., 2020) | DAISIE-DE | 0.080122 | 1.421803 | 0.391713 | 0.313093 | -230.5491 |
|  | DAISIE | 0.080147 | 1.421845 | 0.391762 | 0.313127 | -230.5491 |

To further assess the robustness of DAISIE-DE’s inference, we performed a parametric bootstrap analysis using one of the four empirical datasets, the Galapagos birds dataset. We fitted the model to the Galapagos birds dataset, and we simulated 5000 datasets under the model using the ML estimated parameters from the Galapagos birds dataset. We then re-estimated parameters for each simulated dataset and compared how much these estimates vary across the 5000 replicates. Fig. 3 shows the distribution of parameter estimates across the 5000 replicates. This figure shows that the median values of the estimated rates of cladogenesis, colonization, anagenesis and extinction are close to the true generating values.

**Figure 3.**
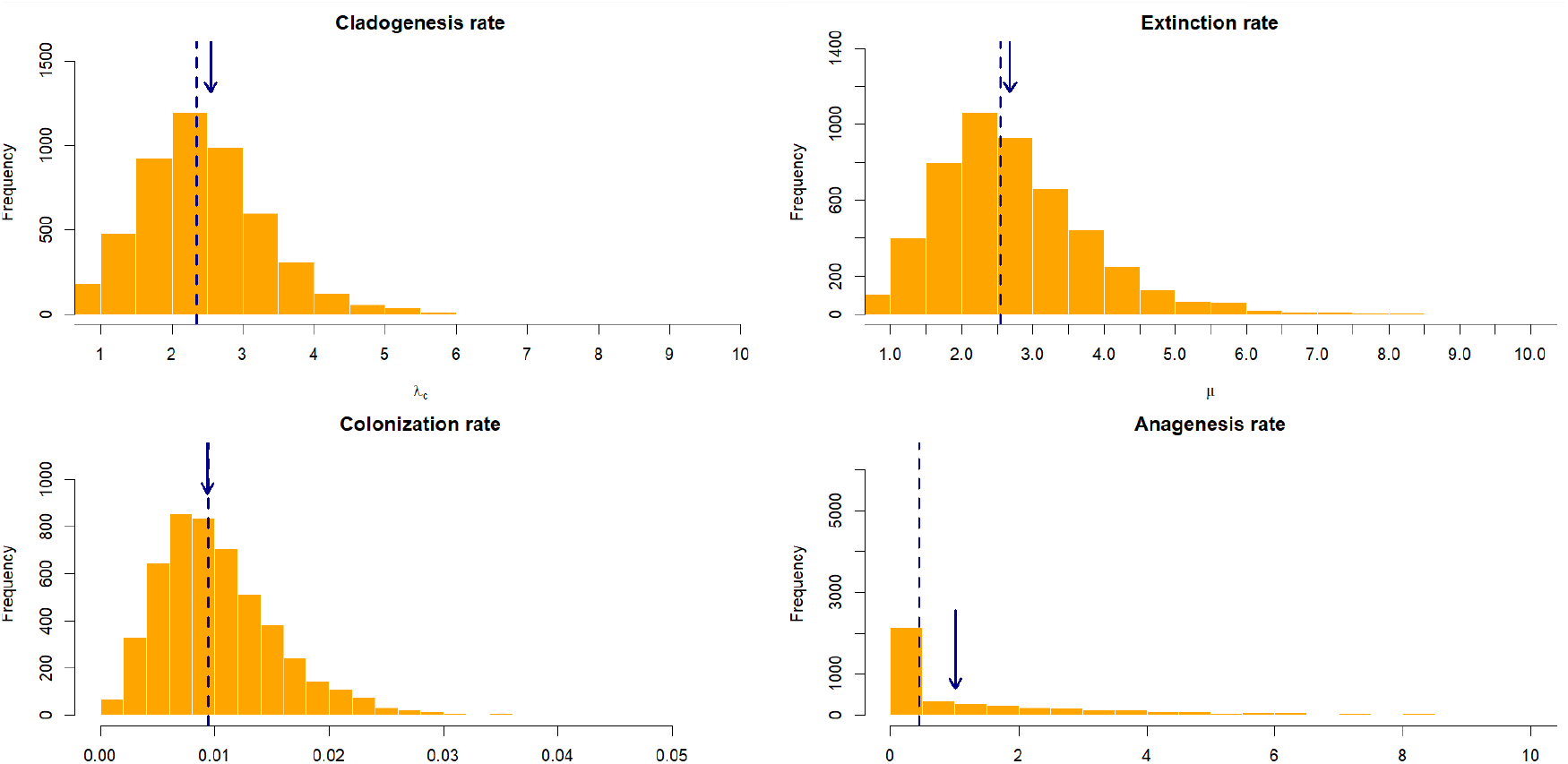
Goodness of fit of the DAISIE-DE model. The arrows indicate the true values of the rates of colonization, cladogenesis, anagenesis and extinction used to generate the simulated datasets. The dashed lines represent the median of the distribution of each parameter estimate. The anagenesis rate is plotted on a logarithmic scale

## 4 Discussion

In this paper, we introduced an alternative approach for computing the likelihood of island communities assembly through colonization, speciation, and extinction. The main motivation behind this reformulation was to enable the incorporation of additional biological complexity such as trait dependence in colonization and diversification rates, which is difficult to implement within the standard DAISIE framework.

To evaluate the performance of our new approach DAISIE-DE, we applied it to four empirical island biota datasets and compared its estimates of colonization, speciation (cladogenesis, anagenesis), and extinction rates with those obtained using the original DAISIE model (see Table 1). We found a strong agreement between the two models across all datasets, indicating that DAISIE-DE preserves the inferential robustness of DAISIE. In addition, DAISIE-DE is also computationally more efficient. The new likelihood calculation requires less computational time and memory than the original DAISIE implementation. This is particularly important as the new diversity-independent model can be applied to much larger trees, making DAISIE-DE a practical alternative in these cases. Of course, one has to bear in mind that it cannot be used when colonization or diversification are diversity-dependent.

More importantly, DAISIE-DE is built upon the same framework as State-dependent Speciation and Extinction (SSE) models, which opens the door to future extensions, such as incorporating trait-dependent diversification and colonization. We will explore this in a future study. Furthermore, the similarity between DAISIE and DAISIE-DE lends support to the validity of the original DAISIE model itself as two fundamentally different frameworks yield identical results across diverse empirical datasets.

## ACKNOWLEDGMENTS

We would like to thank Ryan Brewer, Frederic Lens, Thijs Janzen and Luis Valente for their input, the Center for Information Technology of the University of Groningen for their support and for providing access to the Peregrine and Hábrók high-performance computing clusters, and the Dutch Research Council for financial support through a KLEIN grant awarded to Frederic Lens and Rampal S. Etienne (OCENW.KLEIN.498).

## Appendix A Likelihood of lineages with unknown colonization times

Here we present the likelihood calculation for lineages whose exact colonization time is unknown, and for which only an estimated maximum and/or minimum colonization time are/is available. The dynamics of all cases are summarized in Tables 2 and 3. Fig. 4 provides a schematic representation of the state probabilities *D*_*M*_(τ) and *D*_*A*_(τ) defined in the model.

**Table 2:** Differential equations defined for different cases with given colonization time τ_*C*_ over specific time intervals. gration runs from right to left and the initial conditions in a column connect the equations to the equations to The shading colors mean that the same set of equations is being used.

| Type of lineages | $[\tau_p, \tau_1]$ | $[\tau_1, \tau_0]$ |
| --- | --- | --- |
| No surviving descendants | **Initial conditions:**<br>$D_A^{(1)}(\tau_p) = 1, D_M^{(1)}(\tau_p) = 0, E(\tau_p) = 0$ | |
| | **System of equations:**<br>$\frac{dD_A^{(1)}}{d\tau} = -\gamma D_A^{(1)} + \gamma D_M^{(1)}$ $\frac{dD_M^{(1)}}{d\tau} = -(\lambda^a + \lambda^c + \mu)D_M^{(1)} + (\lambda^a E + \lambda^c E^2 + \mu)D_A^{(1)}$ $\frac{dE}{d\tau} = \mu - (\lambda^c + \mu)E + \lambda^c E^2$ | |
| Non-endemic lineage (colonized at $\tau_1$) | **Initial conditions:**<br>$D_M^{(2)}(\tau_p) = 1, E(\tau_p) = 0$ | **Initial conditions:**<br>$D_A^{(1)}(\tau_1^+) = D_M^{(1)}(\tau_1^+) = \gamma d\tau_1 D_M^{(2)}(\tau_1^-), E(\tau_1^+) = E(\tau_1^-)$ |
| | **System of equations:**<br>$\frac{dD_M^{(2)}}{d\tau} = -(\lambda^a + \lambda^c + \mu + \gamma)D_M^{(2)}$ $\frac{dE}{d\tau} = \mu - (\lambda^c + \mu)E + \lambda^c E^2$ | **System of equations:**<br>$\frac{dD_A^{(1)}}{d\tau} = -\gamma D_A^{(1)} + \gamma D_M^{(1)}$ $\frac{dD_M^{(1)}}{d\tau} = -(\lambda^a + \lambda^c + \mu)D_M^{(1)} + (\lambda^a E + \lambda^c E^2 + \mu)D_A^{(1)}$ $\frac{dE}{d\tau} = \mu - (\lambda^c + \mu)E + \lambda^c E^2$ |
| Endemic singleton lineage (colonized at $t_1$) | **Initial conditions:**<br>$D_E(\tau_p) = 1, D_A^{(3)}(\tau_p) = 1, D_M^{(3)}(\tau_p) = 0,$<br>$D_M^{(2)}(\tau_p) = 0, E(\tau_p) = 0$ | **Initial conditions:**<br>$D_A^{(1)}(\tau_1^+) = D_M^{(1)}(\tau_1^+) = \gamma d\tau_1 D_M^{(2)}(\tau_1^-), E(\tau_1^+) = E(\tau_1^-)$ |
| | **System of equations:**<br>$\frac{dD_E}{d\tau} = -(\lambda^c + \mu)D_E + 2\lambda^c D_E E$ $\frac{dD_M^{(2)}}{d\tau} = -(\lambda^a + \lambda^c + \mu + \gamma)D_M^{(2)} + (\lambda^a D_E + 2\lambda^c D_E E)D_A^{(3)}$ $\frac{dD_M^{(3)}}{d\tau} = -(\lambda^a + \lambda^c + \mu)D_M^{(3)} + (\lambda^a E + \lambda^c E^2 + \mu)D_A^{(3)}$ $\frac{dD_A^{(3)}}{d\tau} = -\gamma D_A^{(3)} + \gamma D_M^{(3)}$ $\frac{dE}{d\tau} = \mu - (\lambda^c + \mu)E + \lambda^c E^2$ | **System of equations:**<br>$\frac{dD_A^{(1)}}{d\tau} = -\gamma D_A^{(1)} + \gamma D_M^{(1)}$ $\frac{dD_M^{(1)}}{d\tau} = -(\lambda^a + \lambda^c + \mu)D_M^{(1)} + (\lambda^a E + \lambda^c E^2 + \mu)D_A^{(1)}$ $\frac{dE}{d\tau} = \mu - (\lambda^c + \mu)E + \lambda^c E^2$ |
| Endemic non-singleton lineage (colonized at $\tau_1$) | **Initial conditions:**<br>$D_E(\tau_p) = 1, D_A^{(3)}(\tau_p) = 1, D_M^{(3)}(\tau_p) = 0,$<br>$D_M^{(2)}(\tau_p) = 0, E(\tau_p) = 0$ | **Initial conditions:**<br>$D_A^{(1)}(\tau_1^+) = D_M^{(1)}(\tau_1^+) = \gamma d\tau_1 D_M^{(2)}(\tau_1^-), E(\tau_1^+) = E(\tau_1^-)$ |
| | **System of equations:**<br>$\frac{dD_E}{d\tau} = -(\lambda^c + \mu)D_E + 2\lambda^c D_E E$ $\frac{dD_M^{(3)}}{d\tau} = -(\lambda^a + \lambda^c + \mu)D_M^{(3)} + (\lambda^a E + \lambda^c E^2 + \mu)D_A^{(3)}$ $\frac{dD_A^{(3)}}{d\tau} = -\gamma D_A^{(3)} + \gamma D_M^{(3)}$ $\frac{dE}{d\tau} = \mu - (\lambda^c + \mu)E + \lambda^c E^2$ | **System of equations:**<br>$\frac{dD_A^{(1)}}{d\tau} = -\gamma D_A^{(1)} + \gamma D_M^{(1)}$ $\frac{dD_M^{(1)}}{d\tau} = -(\lambda^a + \lambda^c + \mu)D_M^{(1)} + (\lambda^a E + \lambda^c E^2 + \mu)D_A^{(1)}$ $\frac{dE}{d\tau} = \mu - (\lambda^c + \mu)E + \lambda^c E^2$ |

**Table 3:**
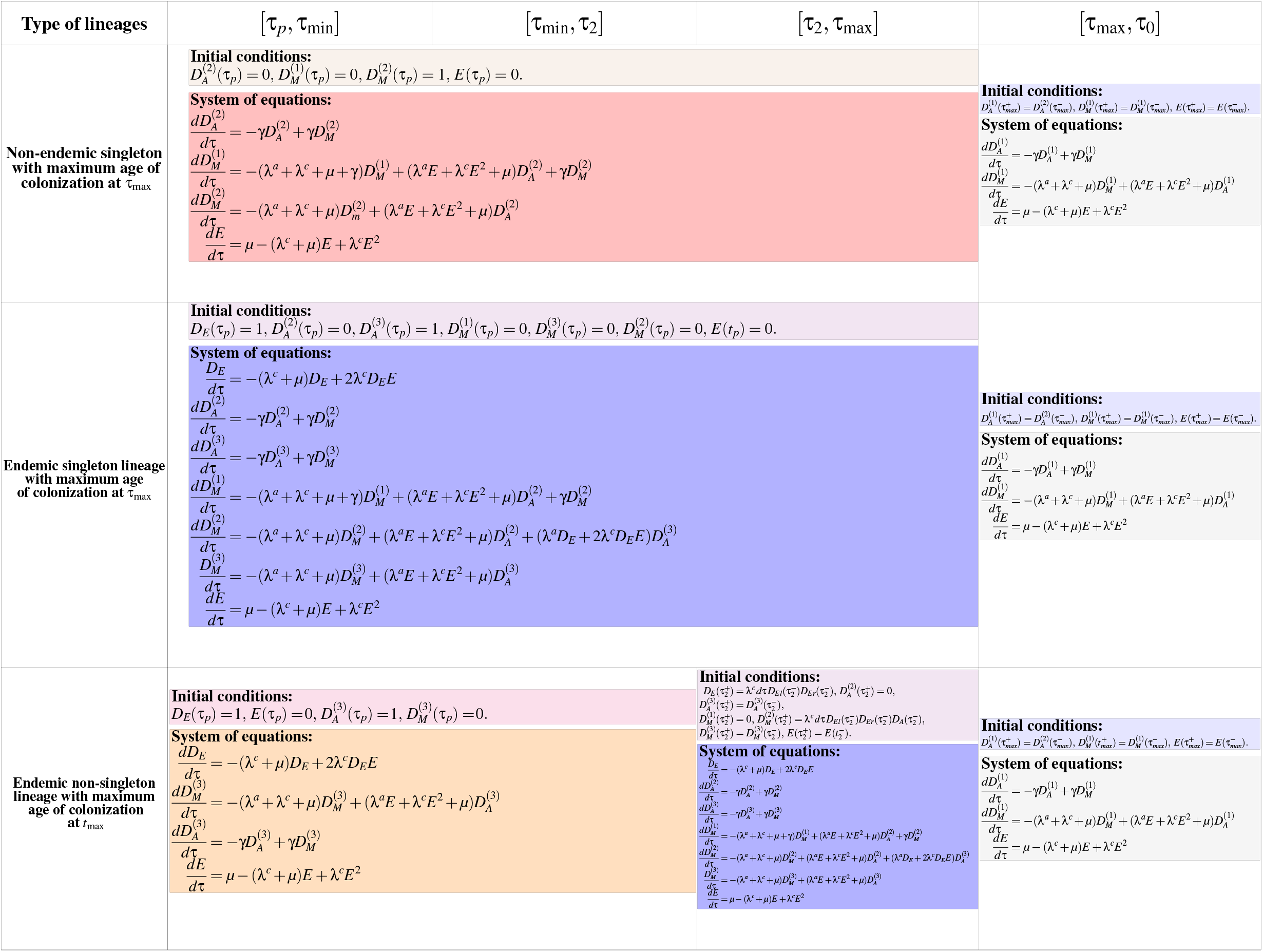
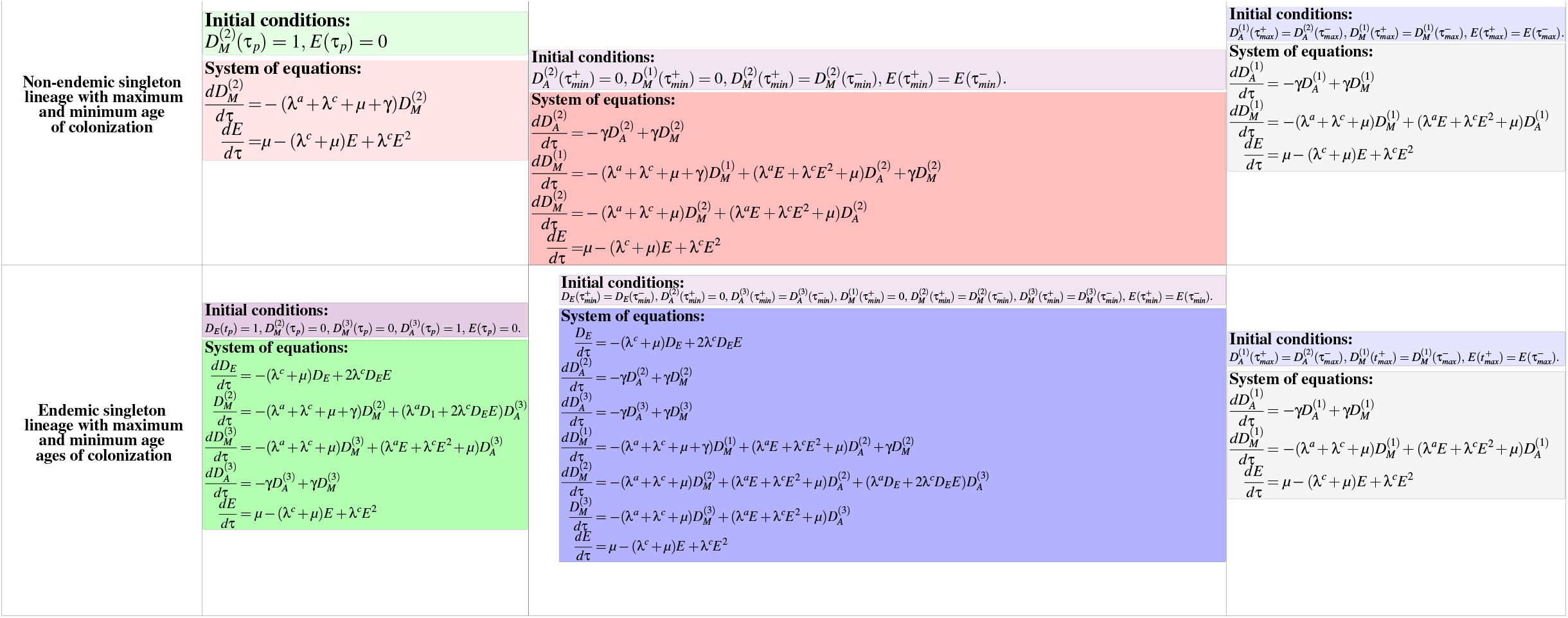
Differential equations defined for different cases with maximum and minimum colonization times over specific time intervals. The integration runs from right to left and the initial conditions in a column connect the equations to the equations to the left. The shading colors mean that the same set of equations is being used.

**Figure 4.**
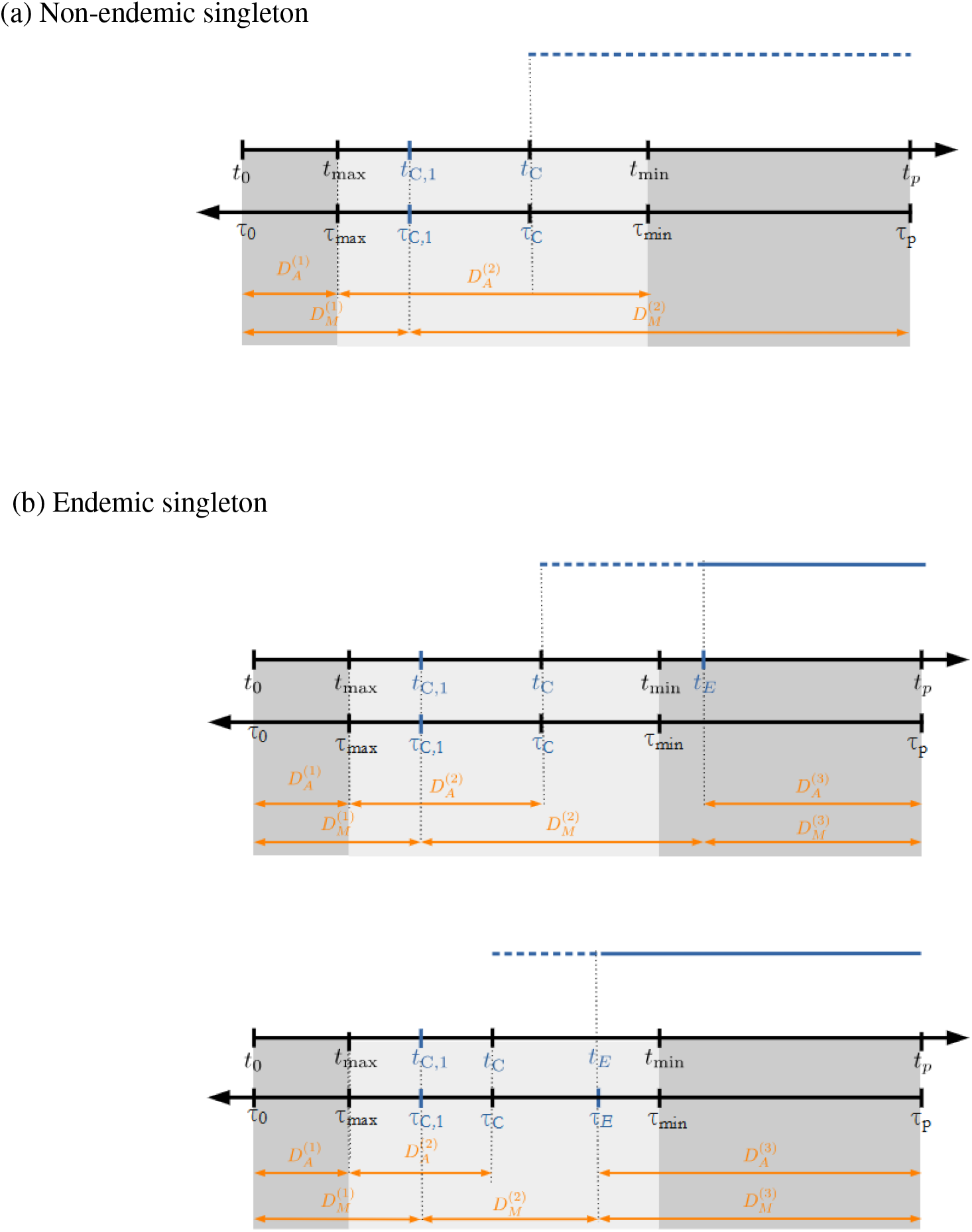
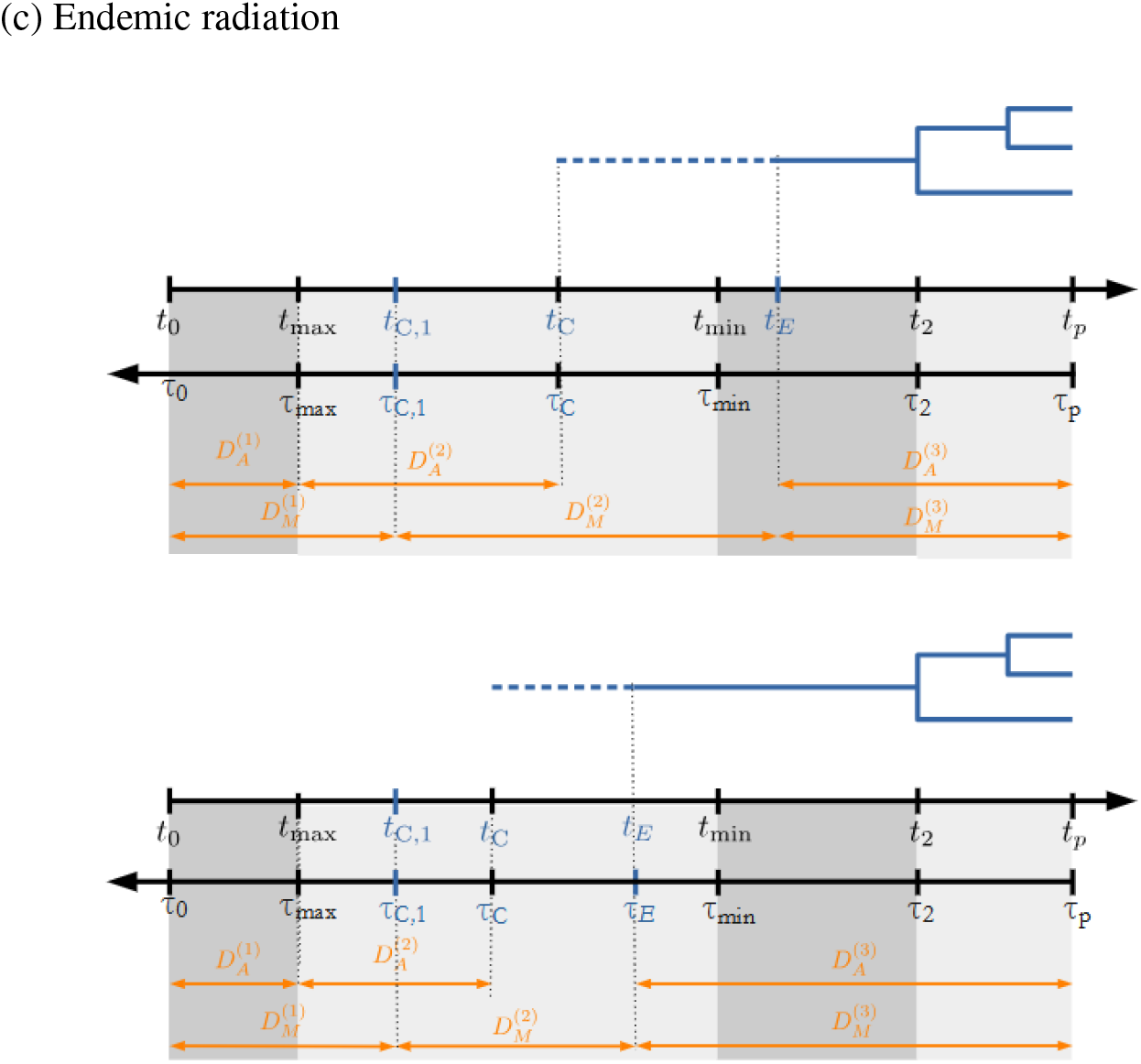
Schematic representations of the temporal structure and definition of the state probabilities *D*_*M*_(τ) and *D*_*A*_(τ) in the backward pruning algorithm for different cases. The blue line represents the time interval during which the species is endemic. The orange arrows show the time intervals over which each probability is computed, going backward in time. The grey-shaded regions represent different time intervals, each governed by a distinct set of differential equations. These intervals correspond to different phases in the lineage history, reflecting changes in model dynamics across time.

The times shown in black (τ_0_, τ_*max*_, τ_*min*_, τ_2_ and τ_*p*_) are extracted from the data. In contrast, the time points shown in blue (τ_*C*,1_, τ_*C*_ and τ_*E*_) are realization-dependent. These times are described as follows:

- *t*_0_, τ_0_: time of the island’s origin, i.e. the island’s age
- *t*_*max*_, τ_*max*_: maximum time of colonization
- *t*_*min*_, τ_*min*_: minimum time of colonization
- *t*_*C*,1_, τ_*C*,1_: time of first colonization event in the colonization time window [*t*_max_,*t*_min_]
- *t*_*C*_, τ_*C*_: time of colonization event that gives rise the lineage observed at the present time
- *t*_*E*_, τ_*E*_ : time at which the lineage at the present time becomes endemic
- *t*_*p*_, τ_*p*_: present day time

There are two possible orderings of these times:

- Upper panels in Figs. 4 b) and c): *t*_*C*_ *< t*_min_ *< t*_*E*_ (including the case *t*_*E*_ = *t*_2_), if speciation of the observed lineages occurs after *t*_*min*_
- Lower panels in Figs. 4 b) and c): *t*_*C*_ *< t*_*E*_ *< t*_min_, if speciation of the observed lineages occurs between [*t*_max_,*t*_min_]

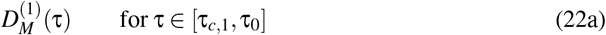

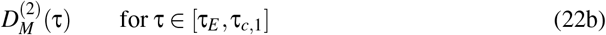

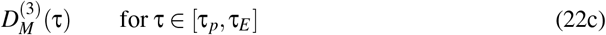

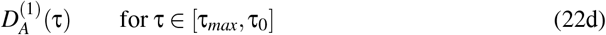

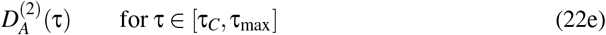

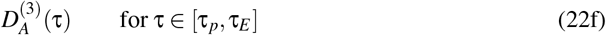

### A.1 Non-endemic lineage with maximum and minimum colonization time

In this case, a non-endemic lineage is present on an island, with known maximum and minimum ages of colonization.

In the interval [τ_*p*_, τ_*min*_], the species is present and is non-endemic. The dynamical equations describing the change in the probabilities in this time interval are given by:

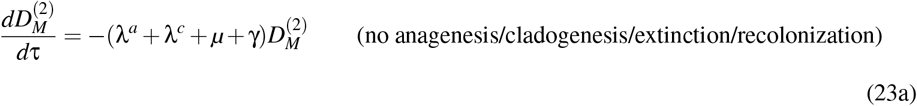

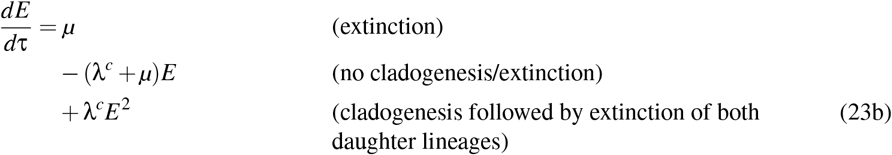

The observed species colonized the island between the maximum colonization time τ_max_, and the minimum colonization time τ_*min*_. If there is a colonization event in [τ_0_, τ_max_] and the mainland species survives until τ_max_, it should be replaced by a new colonization event after τ_max_. To impose this, we keep track in the interval [τ_*min*_, τ_*max*_] of whether the mainland species currently present originates from a colonization before or after τ_*max*_. Hence, we need two variables 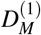 and 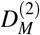. All colonization events in [τ_0_, τ_max_] will be tracked using 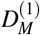. A new colonization after τ_*max*_ corresponds to a transition from 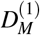 (mainland species present, colonization before τ_*max*_) to 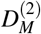 (mainland species present, colonization after τ_*max*_). Therefore, in the interval [τ_*min*_, τ_*max*_] the dynamical equations describing the change in the probabilities are given by:

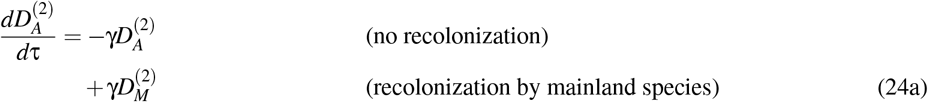

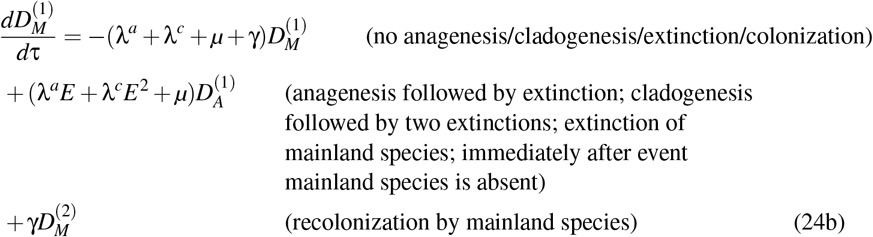

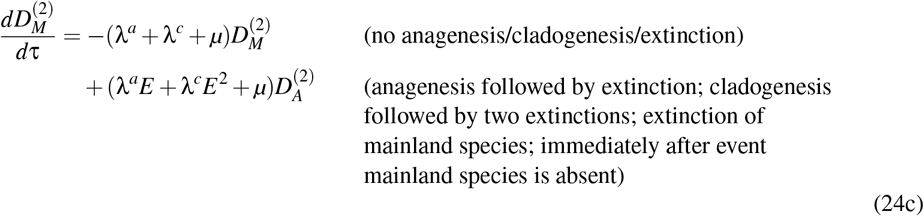

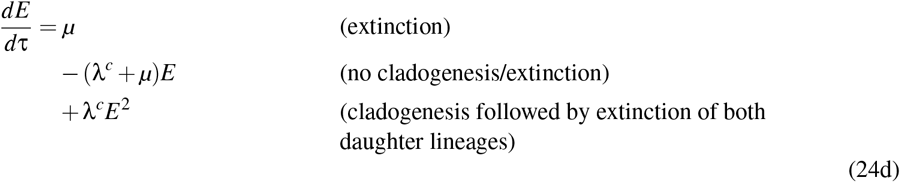

with the initial conditions

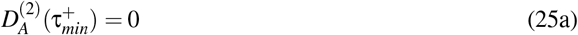

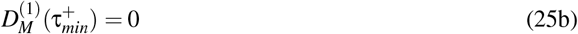

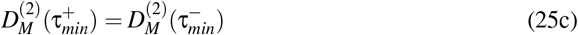

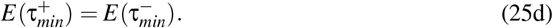

In the interval [τ_*max*_, τ_0_], the dynamical equations describing the change in the probabilities are given by:

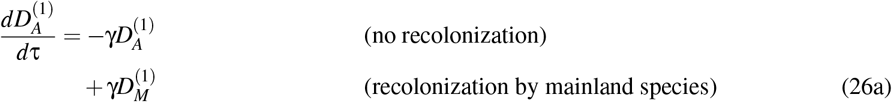

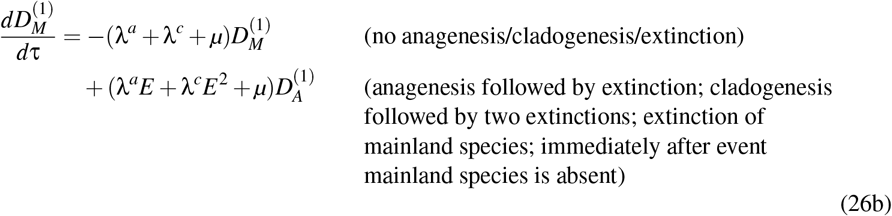

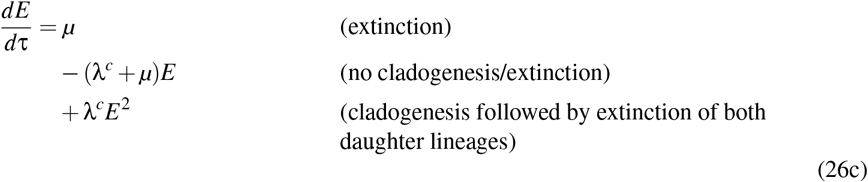

with initial conditions

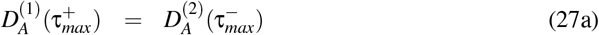

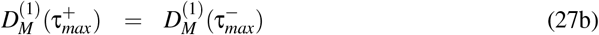

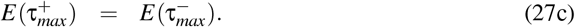

At the island age *t*_0_, we have

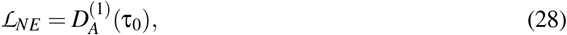

which is the probability that the process that starts at the island age *t*_0_ as an empty island gives rise to a non-endemic lineage that colonizes the island between [*t*_*max*_, *t*_*min*_] and survives until the present time *t*_*p*_.

### A.2 Endemic singleton lineage with maximum and minimum colonization time

In this case, an endemic singleton lineage is present on an island, with known maximum and minimum ages of colonization. If there is a colonization event in [τ_0_, τ_max_] and the mainland species survives until τ_max_, it should be replaced by a new colonization event after τ_max_ (in forward time). To impose this, we should keep track in the interval [τ_*max*_, τ_*min*_] of whether the mainland species currently present originates from a colonization before or after τ_*max*_. Hence, we need two variables 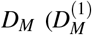 and 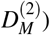. A new colonization after τ_*max*_ corresponds to a transition from 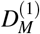 (mainland species present, colonization before τ_*max*_) to 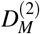 (mainland species present, colonization after τ_*max*_). Between the speciation time and the present the mainland species can recolonize (and go extinct again before τ_*p*_). This possibility is described by the variables 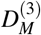 and 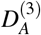.

In the interval [τ_*p*_, τ_*min*_], the species is present, and may either still be non-endemic or have already undergone speciation. The dynamical equations describing the change in the probabilities in this time interval are given by:

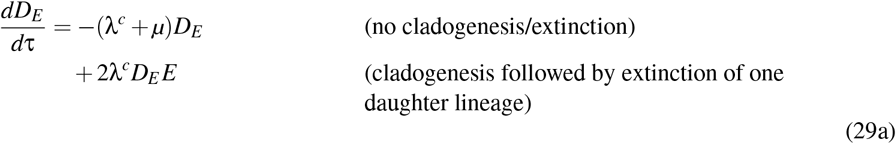

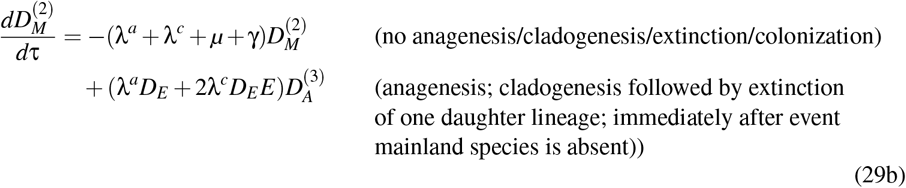

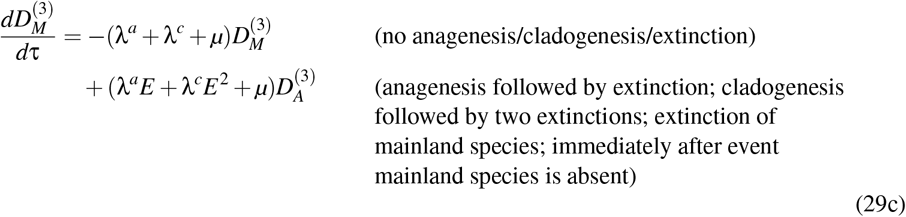

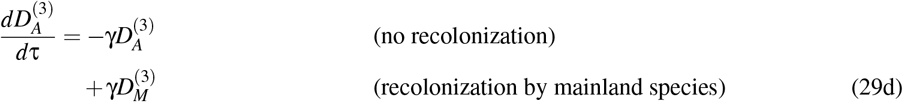

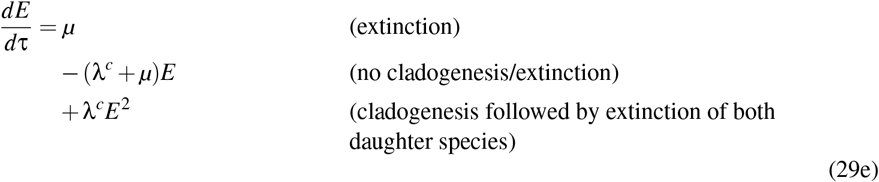

with the initial conditions at τ_*p*_ given by

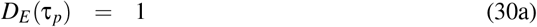

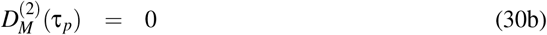

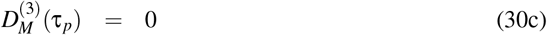

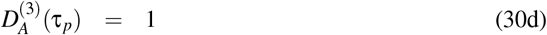

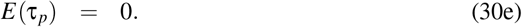

For the interval [τ_*min*_, τ_*max*_], the species has colonized and may or may not have undergone speciation. The dynamical equations describing the change in the probabilities in this time interval are given by:

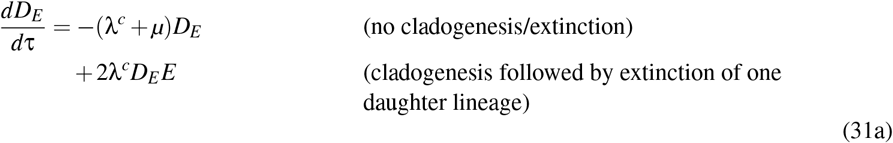

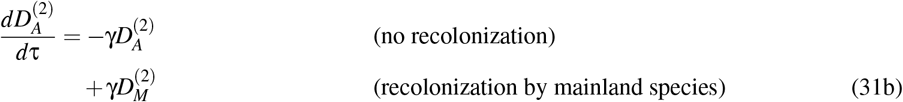

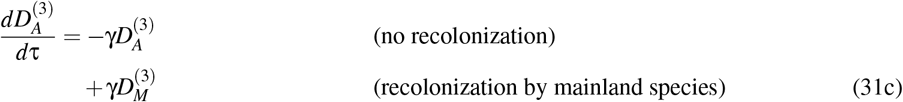

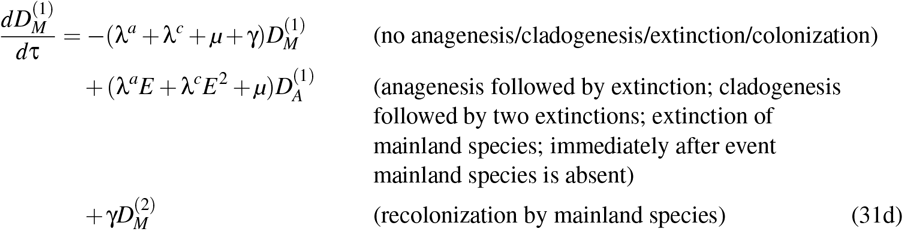

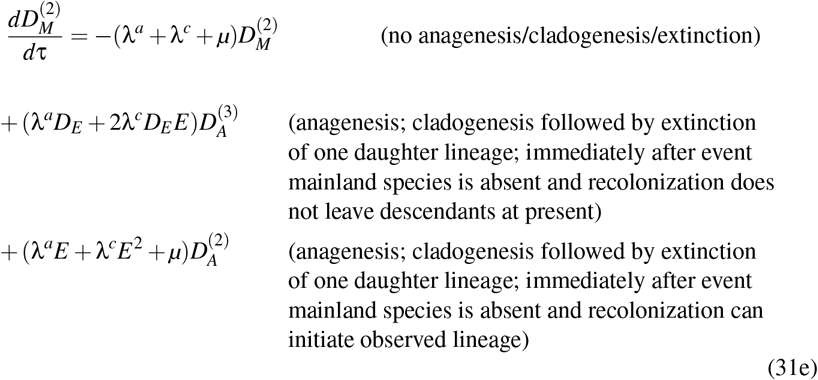

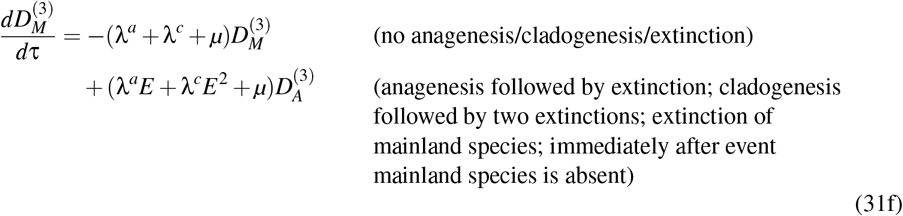

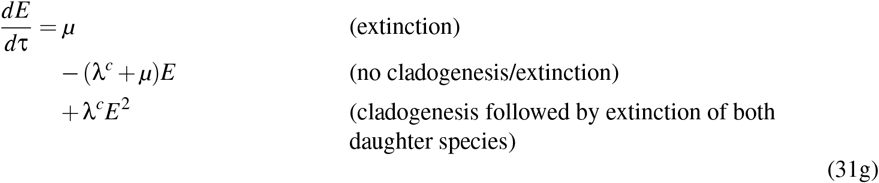

with the initial conditions

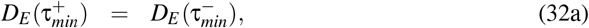

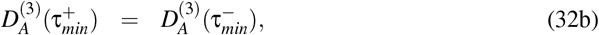

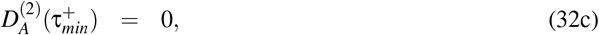

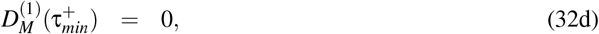

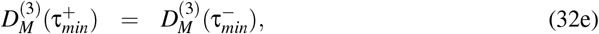

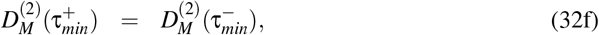

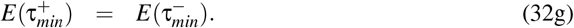

For the interval [τ_*max*_, τ_0_], the dynamical equations are given by eqs. (26a − 26c), with initial conditions

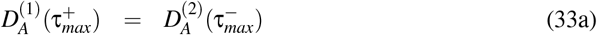

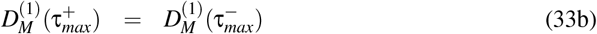

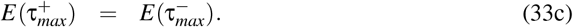

The likelihood is given by

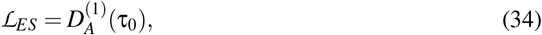

which is the probability that the process that starts at the island age *t*_0_ as an empty island gives rise to an endemic singleton lineage that colonizes the island between [*t*_*max*_, *t*_*min*_], and survives until the present time *t*_*p*_.

If the endemic singleton species coexists on the island with its mainland ancestors, the initial conditions of the set of equations eqs. (29a−29e) are modified to

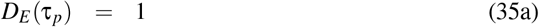

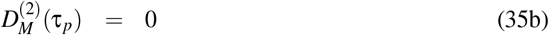

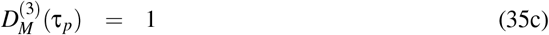

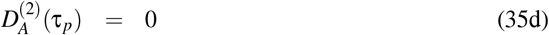

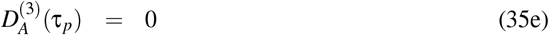

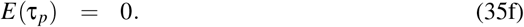

### A.3 Endemic non-singleton lineage with maximum colonization time

In this case, an endemic lineage is present on an island, for which only the maximum age of colonization is known.

In the interval [τ_2_, τ_*p*_], the species has already undergone cladogenesis speciation. For the branches represented in the clade, the dynamical equations describing the change in the probabilities in this time interval are given by:

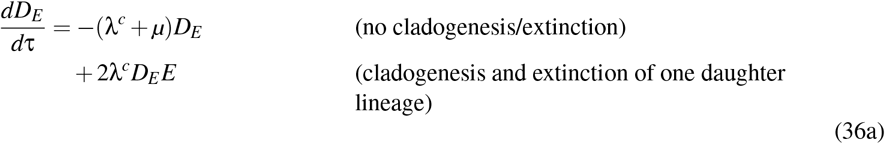

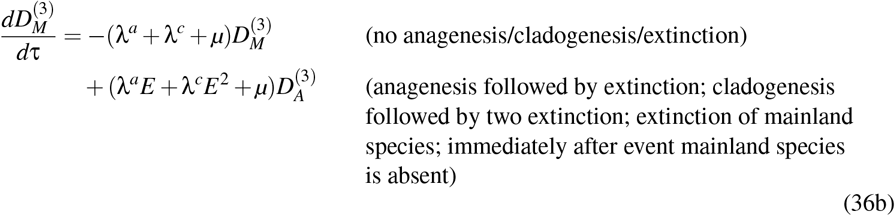

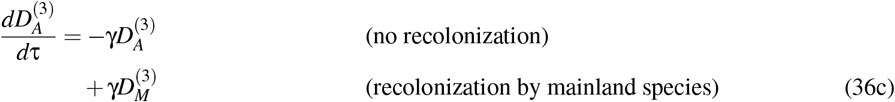

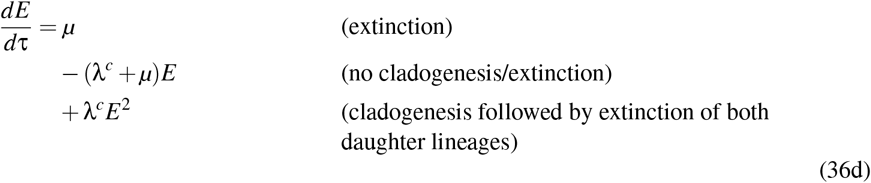

with the initial conditions

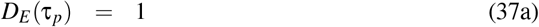

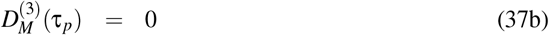

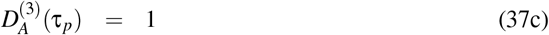

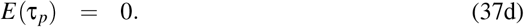

This set of equations is integrated along the edges of the trees starting from the tips. At each branching time we have to stop the integration and set *D*_*E*_ (τ_*i*_) to λ^*c*^*d*τ times the product of the *D*_*E*_ functions of the merging subclades (and do not change the *E* function).

In the time interval [τ_*max*_, τ_2_] we have to add the dynamical equation for 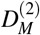, because we know that on at least a part of the branch the species is non-endemic. The dynamical equations describing the change in the probabilities in this time interval are given by eqs. (31a 31g). As initial conditions for 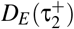 we take λ^*c*^*d*τ times the product of the *D*_*E*_ functions of the merging subclades. For 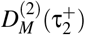 we use the same quantity multiplied by the probability 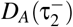 that the mainland species is not present on the island. This covers both cases in which the speciation event that transforms the mainland species into an endemic one occurs either between τ_*max*_ and τ_2_ or precisely at time τ_2_. The initial conditions are then given by:

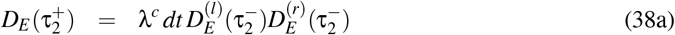

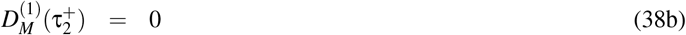

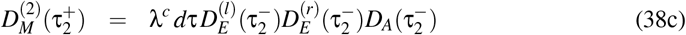

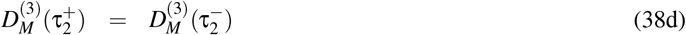

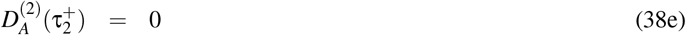

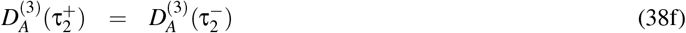

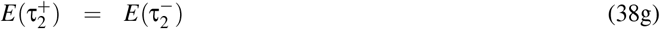

If species within lineages are sampled at random (all species in the clade have the sample probability of being sampled), incomplete sampling can be accounted for by modifying the initial conditions for each tip in the tree to reflect ρ, the probability of sampling a species (following FitzJohn et al. (2009)). In this case, the initial conditions become *D*_*E*_ (τ_*p*_) = ρ and *E*(τ_*p*_) = 1 − ρ.

In the interval [τ_*max*_, τ_0_], the dynamical equations describing the change in the probabilities are given by eqs. (26a−26c) with initial conditions

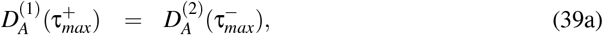

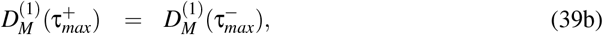

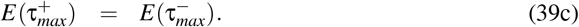

The likelihood is given by

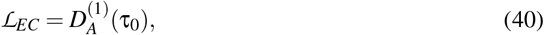

which is the probability that the process that starts at the island age *t*_0_ as an empty island gives rise to an endemic clade that colonizes the island between [*t*_*max*_, *t*_2_], and survives until the present time *t*_*p*_.

If the endemic singleton species coexists on the island with its mainland ancestors, the initial conditions of the set of equations eqs. (36a−36d) are modified to:

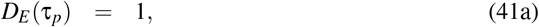

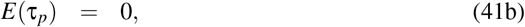

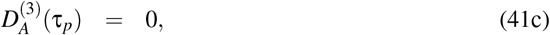

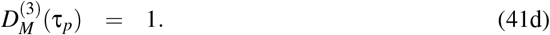

## Appendix B Analytical likelihood expressions for special cases

Here we present analytical likelihood expressions for a series of special cases in order to provide additional support to both the DAISIE and DAISIE-DE models. To facilitate the derivations, we assume that both extinction and anagenesis are absent, i.e., we set *µ* = λ_*a*_ = 0. Let *t*_0_ denote the island age, *t*_*C*_ the colonization time, *t*_2_ the branching time, and *t*_*p*_ the present time. All times are expressed in forward time, so that *t*_0_ *< t*_*C*_ *< t*_2_ *< t*_*p*_.

### B.1 Dynamics before colonization time

Before the colonization event of the observed lineage, the mainland species may have previously colonized the island. If this occurred, the earlier colonizing lineage must not have undergone speciation, as its descendant lineages cannot go extinct under the model assumptions. If a mainland species colonized the island prior to the estimated colonization time of the observed lineage and survive until that time, its dynamics are assumed to be taken over by that of the newly colonizing lineage, in accordance with the DAISIE and DAISIE-DE model assumptions. This gives the following dynamics before the colonization time:

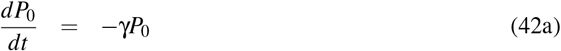

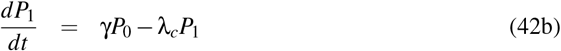

with solutions:

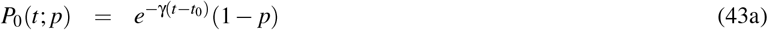

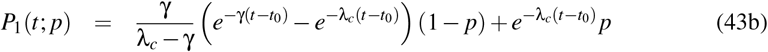

where *p* is the probability that the mainland species is present at time *t*_0_.

The likelihood is given by

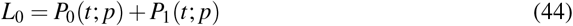

### B.2 Endemic clade with two tips, known colonization time

For an endemic clade that colonizes the island at time *t*_*C*_ and speciated at time *t*_2_, the likelihood formula is:

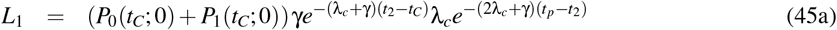

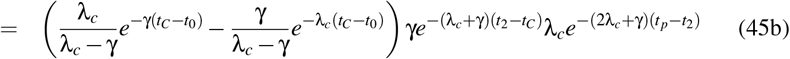

The expression (*P*_0_(*t*_*C*_; 0) + *P*_1_(*t*_*C*_; 0)) describes the probability dynamics before the colonization event at time *t*_*C*_. The exponential term 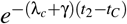 represents the probability of no events (neither cladogenesis, nor extinction) occurring between the colonization time *t*_*C*_ and the cladogenesis event at *t*_2_. Finally, the term 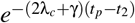 accounts for the probability that both daughter lineages, resulting from cladogenesis at *t*_2_ survive until the present time *t*_*p*_ without further diver-sification and colonization.

### B.3 Endemic clade with two tips, unknown colonization time

The likelihood formula is:

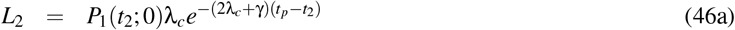

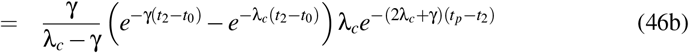

*P*_1_(*t*_2_; 0) describes the probability of observing a singleton lineage on the island, that have colonized at a time *t* between the island age *t*_0_, and the cladogenesis time *t*_2_, the term 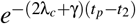 accounts for the probability that both daughter lineages, resulting from cladogenesis at *t*_2_ survive until the present time *t*_*p*_ without further diversification and colonization. It can be checked that:

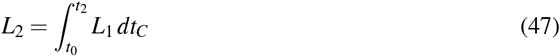

### B.4 Endemic clade with two tips, maximum colonization time

If there is a maximum colonization time *t*_*m*1_, and the branching time *t*_2_ is the minimum colonization time, we subtract from the probability that a colonization takes place before *t*_2_ (*P*_1_(*t*_2_; 0)) the probability that a colonization takes place before *t*_*m*1_ without a new colonization between *t*_*m*1_ and *t*_2_. The likelihood formula is:

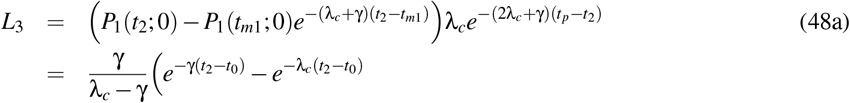

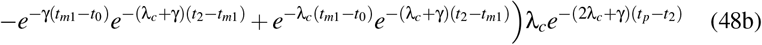

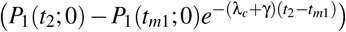 describes the probability of observing a singleton lineage on the island, that have colonized between the time *t*_*m*1_ and the cladogenesis time *t*_2_. The term 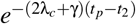 accounts for the probability that both daughter lineages, resulting from cladogenesis at *t*_2_ survive until the present time *t*_*p*_ without further diversification and colonization. *L*_2_ can be recovered from *L*_3_ by setting *t*_*m*1_ = *t*_0_.

### B.5 Endemic clade with two tips that colonized at time *t*_0_

If we know that at time *t*_0_ the island is already colonized by the mainland species, the likelihood would be:

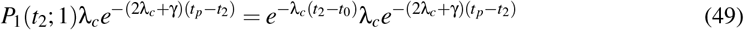

If the mainland species is present on the island with probability *p*, the likelihood is:

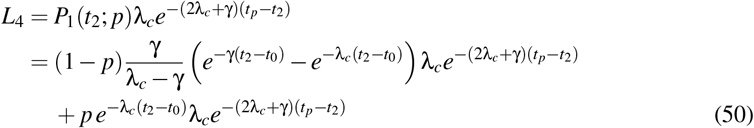

Here, we assume that if the mainland species is not present on the island at time *t*_0_, it can immigrate immediately afterwards.

